# Modeling dynamic allocation of effort in a sequential task using discounting models

**DOI:** 10.1101/839456

**Authors:** Cuevas Rivera Darío, Strobel Alexander, Goschke Thomas, Stefan J. Kiebel

**Affiliations:** Chair of Neuroimaging, Faculty of Psychology. Technische Universität Dresden, 01062 Dresden, Germany; Chair of Differential and Personality Psychology, Faculty of Psychology, Technische Universität Dresden, 01062 Dresden, Germany; Chair of General Psychology, Faculty of Psychology, Technische Universität Dresden, 01062 Dresden, Germany

## Abstract

Most rewards in our lives require effort to obtain them. It is known that effort is seen by humans as carrying an intrinsic disutility which devalues the obtainable reward. Established models for effort discounting account for this by using participant-specific discounting parameters inferred from experiments. These parameters offer only a static glance into the bigger picture of effort exertion. The mechanism underlying the dynamic changes in a participant’s willingness to exert effort is still unclear and an active topic of research. Here, we modeled dynamic effort exertion as a consequence of effort- and probability-discounting mechanisms during goal reaching, sequential behavior. To do this, we developed a novel sequential decision-making task in which participants make binary choices to reach a minimum number of points. Importantly, the time points and circumstances of effort allocation are decided by participants according to their own preferences and not imposed directly by the task. Using the computational model to analyze participants’ choices, we show that the dynamics of effort exertion arise from a combination of changing task needs and forward planning. In other words, the interplay between a participant’s inferred discounting parameters is sufficient to explain the dynamic allocation of effort during goal reaching. Using formal model comparison, we also infer the forward-planning strategy used by participants. The model allows us to characterize a participant’s effort exertion in terms of only a few parameters. Moreover, the model can be adapted to a number of tasks used in establishing the neural underpinnings of forward-planning behavior and meta-control, allowing for the characterization of behavior in terms of model parameters.

## Introduction

It has been known for long that physical effort appears to bear an inherent cost both in humans and other animals (Hull, 1943; Walton, Kennerley, Bannerman, Phillips, & Rushworth, 2006). Although the nature of cognitive effort remains elusive (Shenhav et al., 2017), the role of mental effort has been studied more recently in the same vein (Apps, Grima, Manohar, & Husain, 2015; Kool, McGuire, Rosen, & Botvinick, 2010; Pessiglione, Vinckier, Bouret, Daunizeau, & Le Bouc, 2018; Schmidt, Lebreton, Cléry-Melin, Daunizeau, & Pessiglione, 2012), as well as its neural underpinnings, e.g., (Radulescu, Nagai, & Critchley, 2015). Generally, effort seems to carry a disutility that diminishes the value of reward an action entails, a phenomenon known as effort discounting (Botvinick, Huffstetler, & McGuire, 2009; Westbrook, Kester, & Braver, 2013).

In psychology and economics, much effort has been put into establishing so-called effort discount functions, i.e., parameterized functions of how the subjective value of a reward diminishes as a specific amount of effort is required to obtain it. As with delay- and probability-discounting, several parametric shapes of the effort discounting function have been suggested: hyperbolic (Prévost, Pessiglione, Météreau, Cléry-Melin, & Dreher, 2010), inspired by delay- and probability-discounting; linear (Skvortsova, Palminteri, & Pessiglione, 2014); bilinear (Phillips, Walton, & Jhou, 2007); parabolic (Hartmann, Hager, Tobler, & Kaiser, 2013); and sigmoidal (Klein-Flügge, Kennerley, Saraiva, Penny, & Bestmann, 2015). Additionally, a framework based on prospect theory conceptualizes effort discounting as a shift of the status-quo (Kivetz, 2003). See also (Białaszek, Marcowski, & Ostaszewski, 2017; Klein-Flügge et al., 2015; Talmi & Pine, 2012) for comparisons between these different models.

While these studies established a mathematical description of how required effort affects the valuation of a reward, the experiments are typically constrained to the particular case where the decision to invest effort to obtain reward must be made immediately. However, in most cases of goal-directed behavior in daily life, the reward is not obtainable immediately but must be pursued over an extended time period. This means that in typical effort discounting experiments one cannot address the question of when people will invest effort to obtain a reward that remains obtainable over an extended period of time. For example, an employee may be given a deadline of two weeks to complete an assignment that takes one day. The question for this employee on every day until assignment completion is whether she should invest the effort today or wait until later (Steel & König, 2006). This question is outside the domain of typical effort discounting experiments because there is no ‘wait until later’ option. Some individuals would probably do the assignment early because there may be an unforeseen situation that prevents them from finishing later. Others would prefer to wait and intend to do the assignment late, e.g., just before the deadline runs out, because perhaps it turns out that the assignment is no longer required. Clearly, all possible courses of actions (do the effort early or late) have their advantages and disadvantages and put individuals into a decision dilemma. We believe that this dilemma is central to the meta-control question of how effort discounts potential reward because the dilemma emerges typically when one is pursuing goals that cannot be obtained now but only after some extended time (Goschke, 2014).

In order to induce this dilemma, it is necessary to put participants in a situation where forward planning and future contingencies are important, as opposed to the single-trial experiments traditionally used to elicit discounting. By forward planning, we mean that to make a decision one has to plan several time steps into the future to predict the consequences of possible courses of actions (Dolan & Dayan, 2013). For example, the employee may on day one simulate through in her mind several alternatives of when to do the assignment, select one of these alternatives and execute the first action of this alternative. The question is how one can model decision making in this dilemma by combining forward planning over several trials and previously established effort discounting models for a single trial.

To address this question, we developed a sequential decision making task that captures the effort-investment decision dilemma described above. In each trial of a trial sequence, participants were given the choice to exert effort right away to improve their chances of obtaining a reward at the end of the trial sequence, or wait and not invest effort to see how the situation evolves, so that eventually the need for effort might disappear, however at the price of lowering the chances of reward. We found that the proposed computational model was able to explain different time points at which different participants invested effort. Using formal model comparison, we inferred the forward-planning strategy used by participants during the task. We also show that the inferred effort- and probability-discounting parameters provide for an easily interpretable explanation of the early versus late effort allocation effect observed in the choice data.

In summary, we present a computational-experimental approach, in the form of a novel experimental task and a sequential decision-making model, that enables future studies into the effects of pursuing long-term goals based on moment-by-moment decisions about effort investment in human participants.

## Methods

Participants were recruited from a pool of potential participants organized by the Technische Universität Dresden that includes students as well as individuals from the general population. Of *N* = 60 participants taking part in the experiment, five had to be excluded based on their poor performance during an initial training period. This left *N* = 55 participants (18 female, with an average age of *M* = 26.0, *SD* = 10.8) for our analyses.

Participants went through two different experimental tasks which, together with introduction and training, took an average of 1.5 hours. The two experimental tasks were a single-task effort/probability discounting paradigm and the novel sequential task. In this work, we report only the analysis of the sequential task data that was performed before the single-trial task. For this reason, we describe here only the sequential task.

Payoff was a basic reimbursement of 9 Euros for participating, plus a performance-based bonus of up to 5 Euros for the sequential task. Some participants traded the basic reimbursement for course credit. On average, participants who did not trade the basic reimbursement for course credit earned around 14 Euros for the whole experiment.

The study was approved by the Institutional Review Board of the Technische Universität Dresden (protocol number EK 541122015) and conducted in accordance with the declaration of Helsinki. All subjects gave their written, informed consent.

### Sequential task

In this task, participants are instructed to accumulate points over the course of a mini-block (a trial sequence) of ten trials, with the objective of surpassing a point threshold at the end of a mini-block. To do this, they must, at every trial, choose between a mentally effortful and a probabilistic option, both of which can eventually lead to a reward at the end of a mini-block. At each trial, the current number of points is displayed as a bar shown on the bottom of the screen (see Figure 1B) during the cue and decision phase (see Figure 1C). In order to fill the bar in the mini-block, five points are necessary. If during a mini-block the bar is filled, 20 Euro cents are added to the participant’s final reward. Otherwise, they gain no reward for the mini-block. Each participant went through 25 mini-blocks, plus ten mini-blocks during an initial practice period, for which the participant did not earn any monetary reward. Monetary reward was contingent on winning mini-blocks (as opposed to simply maximizing points) to give special significance to winning a mini-block and to implicitly dissuade participants from focusing on getting the maximum number of points by always choosing the effortful option.

**Figure 1.**
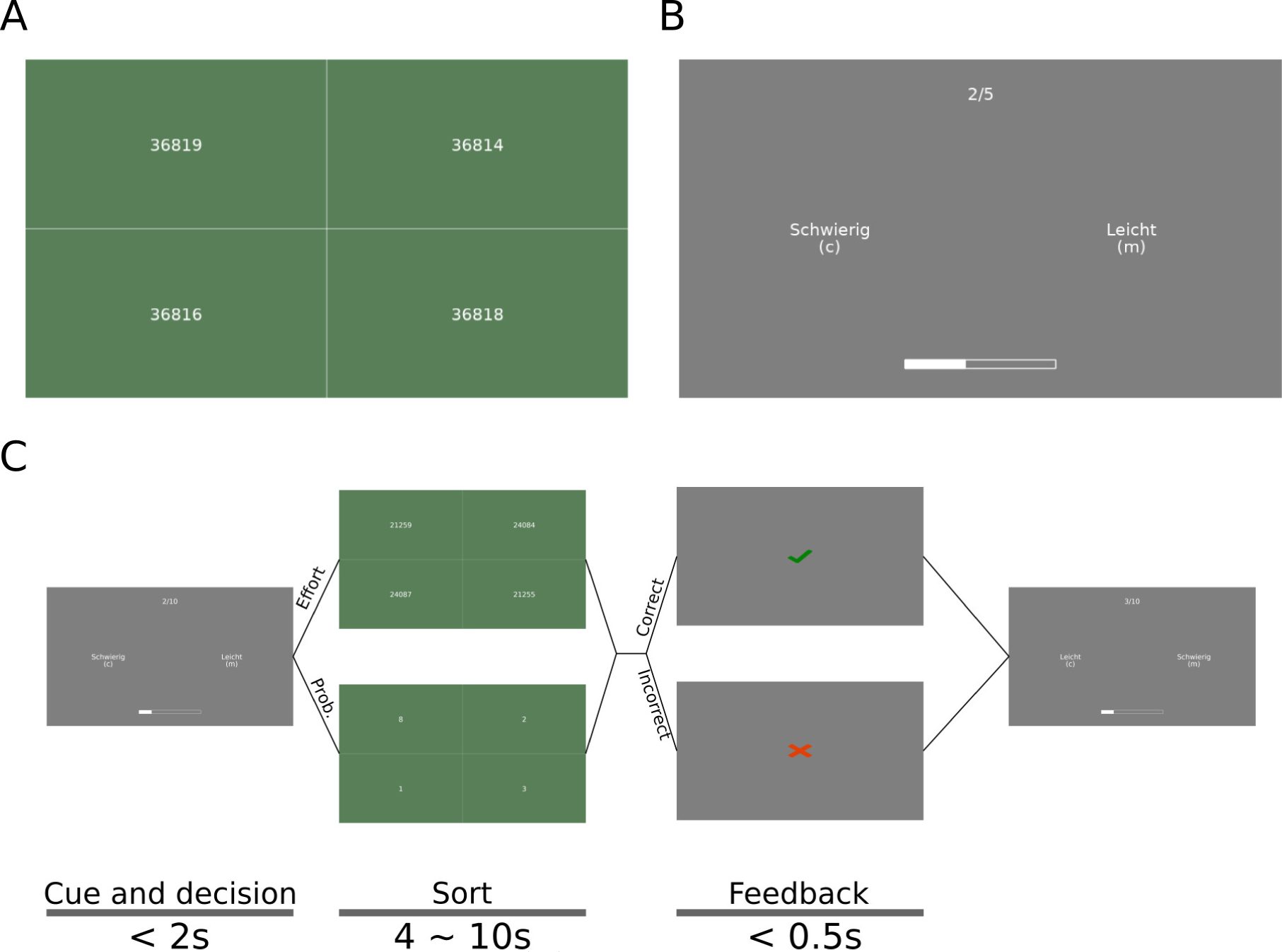
Effort and sequential task. **(A)** Number-sorting task, where participants must select the shown numbers in ascending order to correctly complete a single trial of the sequential effort-investment task. To select a single number, participants can click anywhere in the box containing this number. **(B)** Cue and decision phase of the sequential task. Participants must choose between the easy option (Leicht, in the original German), which corresponds to the probabilistic option (see main text), and a hard option (Schwierig), which corresponds to the effortful option, which leads to the task shown in (A). The choice is made with keyboard keys C and M, for the option on the left and right, respectively; the side on which each option appears is randomly selected at every trial. **(C)** Schedule of the different phases of a single trial in the sequential task. The times for each screen are shown at the bottom, along with the name of each phase. The main experiment consisted of 25 mini-blocks (sequences of trials) with ten trials each.

For the effortful option, we used a number-sorting task, in which a set of numbers is shown on screen with five digits each that can differ in any of the digits (see Figure 1A). The participant must sort the set of numbers in ascending order by sequential mouse clicks on the displayed numbers within a fixed time period. This time is adapted to each participant during training such that their performance on the number-sorting task is around 90% (see Section ‘Procedure’) to equalize the required effort across all participants. If the numbers are sorted correctly, participants gain one point.

For the probabilistic option, participants were to complete a number-sorting task as well, but all numbers had a single digit, rendering the task practically cognitively effortless. Participants had a 50% chance of gaining a point for correctly completing the sorting task, which was told to them in the instructions. The probabilistic option corresponds to waiting until a later trial to exert effort, if it ever becomes necessary. The probability associated with the probabilistic option is included to create mini-blocks in which the participant can win without having to exert any effort by choosing this option at every trial and being “lucky” with the outcomes. We included the single-digit sorting trial to equalize the physical effort that comes from using the mouse to click on the numbers.

Thus, each trial of the sequential task is divided into three phases: (1) the cue and decision phase (Figure 1B), in which participants must choose between the two options using the keyboard (“c” for the option shown on the left, “m” for the option shown on the right). The left/right position of the two options (probabilistic and effortful) on the screen is randomized every trial. This phase lasts until the participant makes the decision, but no longer than three seconds; (2) the sorting phase, in which participants must carry out the selected task. This phase lasts between four and ten seconds, depending on the participant’s performance during training (see below); and (3) the feedback phase, in which participants are told whether they correctly completed the task or not. This phase lasts half a second. Figure 1C shows a diagram of the trial timing, including all the screens observed by participants as well as the timings of each phase of the trials.

Importantly, the number of points required to win a mini-block is only half of the number of trials in the mini-block. This, combined with the 50% chance of getting a point with the probabilistic option, has the effect that, by just choosing the probabilistic option, the participant can win on average half the mini-blocks in the experiment. Additionally, because the difficulty of the effortful task was set such that expected performance is close to 100%, the participant is almost guaranteed to win every mini-block, regardless of the strategy chosen, as long she is willing to invest the effort associated with the effortful option when it becomes necessary, i.e., when she would otherwise risk not having enough points at the end of the mini-block.

### Procedure

The experimental session began with instructions shown on the screen. No instructions were given by the experimenter. Then, the participant went through an introduction to the number-sorting task with the intention of getting them acquainted with how the mouse is used to sort the numbers. During this familiarization period, participants completed twelve trials, divided into six single-digit sorting tasks and six five-digit sorting tasks. Training followed, during which participants’ response times for the main experiment were adjusted. Participants first had to go through a block of 40 trials, in which they had to sort the four numbers as quickly as possible within a fixed time-interval of twelve seconds per trial. This is long enough that no participant timed out. After this initial block, the new interval was chosen to be the 95% percentile of the participant’s reaction times. After that, three more blocks of 40 trials were possible; after each of them, the participant’s performance (i.e., the percentage of times they correctly sorted the numbers before the deadline) was measured. If the performance was below 85%, the deadline was increased. If above 95%, the deadline was decreased. This was repeated for a maximum of four training blocks. If after the training phase the performance was not between 85% and 95%, we excluded the participant from further analysis. The duration of the training phase varied across participants. Once training was done, participants received instructions for the sequential experiment, followed by ten practice mini-blocks, in which they earned no reward (stated in the instructions). Once they finished these, they performed the main experiment with 25 mini-blocks, earning monetary reward for each one completed successfully.

### Single-trial discounting models

The sequential decision-making model proposed in this work is based on classical single-trial discounting models. For completeness, we briefly describe their mathematical form in this section.

It is now well accepted that the best-fitting discounting function for probability discounting is a hyperbola-like one (Ostaszewski, Green, & Myerson, 1998), whose mathematical form is given by:

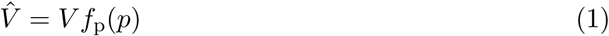

where 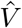 is the subjective value, *p* is the probability of obtaining the reward, V is the objective reward value (e.g. the amount of money) and *f*_p_ is given by:

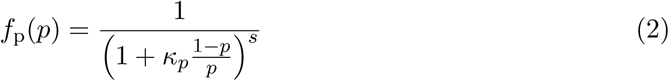

where *κ*_*p*_ and *s* are the model’s free parameters which are to be fit to behavioral data. These two parameters have the effect of creating steeper discounting the higher their values are; *κ*_*p*_ is regarded as a probability-scaling parameter, while *s* is regarded as a non-linear sensitivity to probability (Green & Myerson, 2004).

We made use of this model during our study with one caveat: while the inclusion of the parameter *s* has been previously found to add explanatory power to the model, it makes comparison between groups more difficult (McKerchar & Renda, 2012), as discounting is affected by these two parameters, and it severely complicates parameter fitting due to the high correlation between the parameters (Myerson, Green, & Warusawitharana, 2001). For this reason, we chose to fix *s* to 1 for all participants.

For effort discounting it is less clear which discounting function describes behavioral data best (Białaszek et al., 2017; Kivetz, 2003; Klein-Flügge, Kennerley, Friston, & Bestmann, 2016; Klein-Flügge et al., 2015; Kool et al., 2010). Formal model comparison has been performed between different discount functions, with differing results (Białaszek et al., 2017; Klein-Flügge et al., 2015).

In this work, we exemplify our model using hyperbolic and sigmoid effort discount functions. We chose hyperbolic discounting for its long tradition in probability- and delay-discounting, which makes it a prime candidate for effort discounting. Sigmoid discounting, on the other hand, has the property of being concave for low effort levels and convex for high effort levels, which Klein-Flügge et al. (2015) argued was an integral part of effort discounting. However, note that our modeling approach presented below can be applied to any other discount function.

The hyperbolic effort discount function is given by:

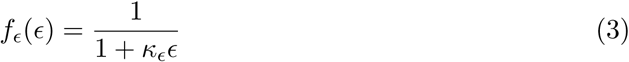

where *ϵ* is the effort level and *κ*_*ϵ*_ is the only free parameter, which, as with probability discounting (Equation 2), represents effort scaling.

The sigmoid discount function is given by:

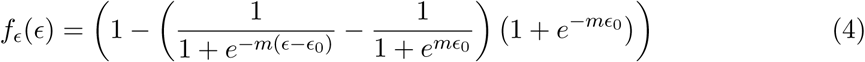

with free parameters *m* and *ϵ*_0_ that correspond to slope of the function at the center (where the value of the function is 0.5) and the coordinate of the center.

While the interpretation of *ϵ* = 0 is clear (there is no effort), effort does not have a natural scale like those of delay and probability. Instead, we chose the units of effort such that the effort level of one number-sorting task is *M* − 1, where *M* is the number of digits of each number to sort. In this scale, the probabilistic option (see Section ‘Sequential task’) has an effort level of zero and the effortful task has an effort level of four.

### Sequential discounting models

In this work we present a novel family of models that bring the single-trial discounting models of the previous section into the realm of sequential decision-making models of goal-directed behavior. To do this, we built on eqs. (1) to (4) and added a component that implements forward planning over future trials to achieve the goal of filling the point bar during a mini-block.

#### Action sequences

For our forward-planning model, we first introduce the concept of action sequences *π*, which we define as a list of actions to perform in future trials, one for every trial left in the mini-block. Because in the sequential task, the participant must make forced choices between an effortful and a probabilistic option, an action sequence consists of these binary choices, one for each remaining trial until the end of a mini-block. For example, at the very beginning of a mini-block (with ten trials left), an action sequence could consist of only the probabilistic choices at every trial in the future. This would be the policy of a participant who, at the beginning of the mini-block, prefers not to choose the effortful options throughout the mini-block. Another would be an action sequence consisting only of choosing the effortful options. Planning for more nuanced strategies is also possible, i.e. a mix of both options.

The model evaluates every possible action sequence in a way that reflects the overarching goal leading to reward, i.e., filling the point bar. Since at every trial the choice is binary, the total number of possible action sequences at the beginning of trial *t* is 2^*T −t*+1^, including the one to be made at trial *t*, where *T* is the total number of trials in a mini-block (ten in our experiment).

It is unlikely that human participants use such a brute-force, binary-tree search algorithm to find the best strategy, as the number of action sequences grows exponentially with the number of trials left; therefore, we created a model in which the only two strategies available are (1) committing to choosing the probabilistic option for the remaining trials in the mini-block and (2) committing to choosing the effortful option for the rest of the mini-block, or until the point bar has been filled. Using only these two action sequences captures the essence of the task, in which a frugal decision-making agent would choose to exert no effort unless it becomes absolutely necessary, and a more reward-sensitive agent (i.e., one that wants to maximize the probability of obtaining reward, disregarding the cost of effort) would prefer exerting effort until the probability of winning the mini-block is high enough to risk the probabilistic option. We discuss the validity and usefulness of this reduction in the number of policies in the Discussion section.

We define these two action sequences with *π*_p_ as the action sequence of all-probabilistic choices and *π*_*ϵ*_ as the action sequence of all-effortful choices. With these, we define the set *A* = {*π*_*p*_, *π*_*ϵ*_}.

For every action sequence *π* ∈ *A* the model must produce an evaluation *z*(*π*) which determines how beneficial this action sequence is for achieving the goal. Then, the model will select an action (probabilistic or effortful) using these valuations. Concretely, the action *a*_*t*_ at trial *t* is sampled according to:

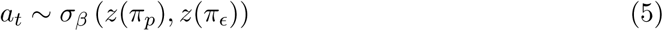

where *σ*_*β*_ is the softmax function with inverse-temperature parameter *β*. We fix the value of this parameter to 5 for all models and participants, which produced posterior probabilities (for effort and probability) in the full range of 0 to 1.

The evaluation function *z* is defined in terms of the single-trial discounting models discussed in Section ‘Single-trial discounting models’. In what follows, we discuss *z*(*π*_*p*_) and *z*(*π*_*ϵ*_) separately.

#### Forward planning with probability

When planning to choose the probabilistic option for every trial into the future, we propose two natural ways of calculating *z*(*π*_*p*_), where one aim of the study will be to use model comparison to disambiguate between these two ways. The first way is to stack the discounting function as many times as there are trials left:

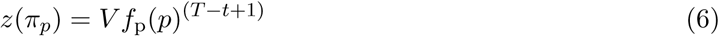

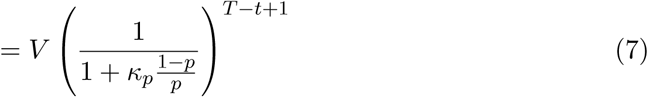

where *f*_p_(*p*) is given by Equation 2 with *s* = 1. This simply means that the objective reward *V* obtained at the end of the mini-block is discounted once for each remaining trial. We refer to this variant as “stack”. Note that we explicitly do not call this variant ‘multiply’ because some other discounting functions (not considered in this paper) are not multiplicative.

With the second variant, one calculates the overall probability of winning the reward by choosing the probabilistic option in every remaining trial in the mini-block, as if it were a single action with an overall probability. The calculation of this overall probability is done with the binomial distribution and the resulting probability is used to apply hyperbolic probability discounting:

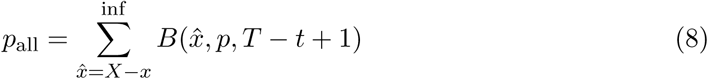

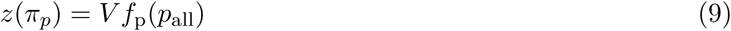

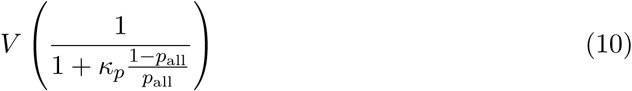

where 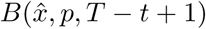 is the probability mass function of the binomial distribution, 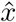 is the number of successes for the binomial, *p* is the probability of success and *T − t* + 1 is the number of trials left; *X* is the number of points necessary to win the mini-block and *x* is the current number of points. *f*_p_(·) is given by Equation 2. We refer to this variant as “add”.

To summarize, with the stack and add variants, we describe two ways to compute at trial *t* the subjective value of reward obtained at the end of the mini-block. The stack variant simply assumes that the single-trial probabilistic discounting is applied as many times as there are remaining trials. The add variant calculates values of probability and effort that take into account the structure of the sequential task and applies the single-trial discount function to these values just once.

#### Forward planning with effort

In analogy to the probabilistic action sequence, we propose two variants of the effortful action sequence evaluation. The first variant is the direct counterpart of the stack variant in probability:

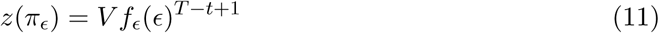

where *f*(*ϵ*) can be hyperbolic effort discounting (Equation 3) or sigmoid effort discounting (Equation 4). As for the probabilistic action sequence, we refer to this version as ‘stack’.

The second variant is the direct counterpart of the add variant in probability, and is defined by adding all the future efforts as if it were a single action and discounting the resulting added effort using the hyperbolic or sigmoid functions:

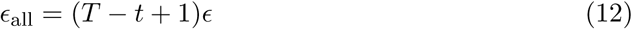

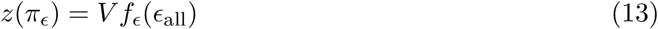

where *f*_*ϵ*_(·) can be the hyperbolic effort discounting (Equation 3) or sigmoid effort discounting (Equation 4). As for the probabilistic action sequence, we refer to this version as ‘add’.

#### Model variants

We define the different variants of the sequential model depending on the type of forward planning used for effort and probability, each of which can be “stack” or “add”. This gives us a total of four variants of the sequential component, naming the effort variant first: add/add, stack/add, add/stack, stack/stack. For example, we refer to the variant in which effort is stacked and probability is added as stack/add.

### Model comparison

In total, we propose a family of eight (2 × 2 × 2) models: (sigmoid or hyperbolic) × (stacking or adding probability) × (stacking or adding effort). In order to select the one that fits our data best, we implemented the hierarchical model proposed by Stephan, Penny, Daunizeau, Moran, and Friston (2009), which we only briefly describe here. Note that Stephan et al. (2009) suggest using the so-called exceedance probability to produce a ranking between several models, which takes into account both how many times each model was inferred to be the best for participants, and the uncertainty derived from the inference procedure, making it a more appropriate measure for model comparison than approximations to the model evidence such as the Bayesian information criterion (Schwarz, 1978).

Stephan et al. (2009) defined a hierarchical model in which the models to be compared are first fit to the data of each participant using Bayesian methods. From this fitting, the model evidence can be calculated for every combination of participant and model. This matrix of model evidences is then used as “data” for the hierarchical model. Formally, the model evidence is introduced as *p*(*d*|*m*), where *d* is the data (participants’ choices) and *m* represents one of the 8 variants we propose, defined as a vector of zeros with a single 1 in the place of the model (for example, the third model is represented by *m* = (0, 0, 1, 0, 0, 0, 0, 0)). This is used to infer, using Bayes theorem, which model best fits the data of all participants together.

The hierarchical model then defines the probability of the model *m* given an auxiliary variable *r*:

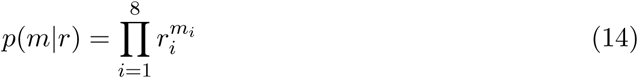

The variable *r*_*i*_ can be interpreted as the number of participants for which model *m*_*i*_ was the best model (highest model evidence), although this is a simplification. The last component to define is the prior probability of *r*, which we defined as a flat Dirichlet distribution (as was done by Stephan et al. (2009) in their examples):

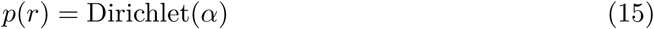

where *α* is a vector of ones, which reflects that we do not have any hypothesis *a priori* regarding which of the variants of our model fits the data best.

Finally, the full generative model is given by:

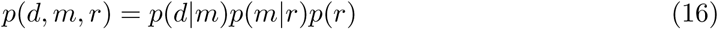

which we inverted to produce the posterior probability *q*(*m*|*d*) by using the NUTS sampler as implemented in PYMC3 (Salvatier, Wiecki, & Fonnesbeck, 2016). These posterior distributions can then be used to perform model comparison via the computation of the exceedance probability, which is a way of determine how much more likely is one model to better describe the data than all other models (Stephan et al., 2009).

To calculate the exceedance probability for model *i*, it suffices to calculate the cummulative distribution of *p*(*r*_*i*_|data) over all values for which *p*(*r*_*i*_|data) > *p*(*r*_*j*_|data), for all *j* ≠ *i*.

### Dividing participants into groups

We divided participants into three groups based on their effort exertion strategy which we determined given their choice data. The first group, called all-effort group, consisted of those participants who chose the effortful option in more than 90% of trials. This implies that these participants used the effortful option even after winning the mini-block.

The remaining participants were divided into two groups: those who applied effort early in the mini-block (early-effort group) and those who applied it late (late-effort group). To divide participants we made use of the frequency of effort calculated at every trial number across mini-blocks. Intuitively, the frequency of effort for participants in the early-effort group decreases as the trial number increases (until the mini-block has been won), while late-effort group increases their frequency with trial number. To quantify this, we calculated the change in frequency of effort between each trial and the next one:

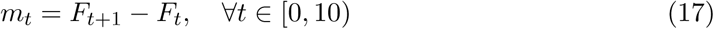

where *F*_*t*_ is the overall frequency of effort for trial number *t*. We found that to classify participants based on when they exerted effort, the best strategy was to count the number of times, for each participant, that the slope was positive for all trials and subtracted the number of times it was negative:

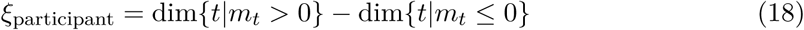

where dim() is a function that returns the number of elements in a set. *ξ*_participant_ determines whether a participant belongs to the early-effort group (*ξ* ≤ 0) or to the late-effort group (*ξ* > 0).

### Parameter estimation

Parameter estimation was done using a variational inference scheme implemented in PYMC3 (Salvatier et al., 2016), which uses the mean-field approximation. The outcome of this Bayesian inference scheme is estimations for the mean and standard deviations of Gaussian posteriors for each model parameter (see Section ‘Sequential discounting models’), providing both a single-point estimate, e.g. the mean of the Gaussian posterior, and estimations for the uncertainty of the inference.

Additionally, the model evidence for all models and participants is calculated as the negative loss produced by PYMC3, which is used for model comparison in Section ‘Model comparison’.

Parameter estimation was done using the following generative model:

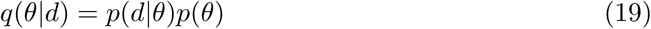

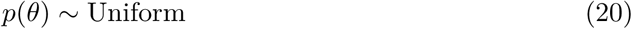

where *p*(·) is a probability distribution, *θ* is the set of parameters to fit to the data and *q*(*θ*) is the posterior distribution over the parameters. Uniform refers to uninformative priors, i.e. prior distributions in which no special prior information is encoded. *p*(*d*|*θ*) is the likelihood function provided by our decision-making model.

## Results

We first show that there are inter-participant differences in the strategies used to reach the goal, which are reflected in the circumstances under which participants chose the effortful option instead of the probabilistic one. Furthermore, we divided the participants according to three behavioral categories, based on their strategies. This is followed by formal Bayesian model comparison to identify the best among eight different models, which differ in terms of how forward planning computes the subjective value of reward, and which out of two discount functions is used. Having selected the best model for our data, we will show that this model correctly captured the overall preference for effort shown by participants. Finally, we show that the overall preference for effort can be understood in terms of the inferred discounting parameters (more specifically, their ratio), providing for an intuitive description of apparent effort preference in participants.

### Behavioral analysis

5 participants were excluded from analysis due to their low success rates in the number-sorting task throughout the experiment. The remaining 55 participants were used for the following analyses.

#### Preference for effort

As a first step to determine whether our task elicited differences in the adaptation of effortful choices between participants, we calculated the overall frequency of effort for each participant in the sequential task, i.e. in which percentage of trials the participant chose the effortful option. The results are summarized in Figure 2A; to determine whether participants had fully understood the instructions regarding reward contingencies (i.e. that gaining points after filling the point bar brings no further reward), the trials were separated into before and after having won the mini-block (i.e. filled the points bar), displayed as blue and green bars, respectively. It can be seen that, on average, participants chose the effortful option much less frequently after having won the mini-block, which is congruent with the rules of the task (i.e. that getting more points after having filled the bar is of no use).

**Figure 2.**
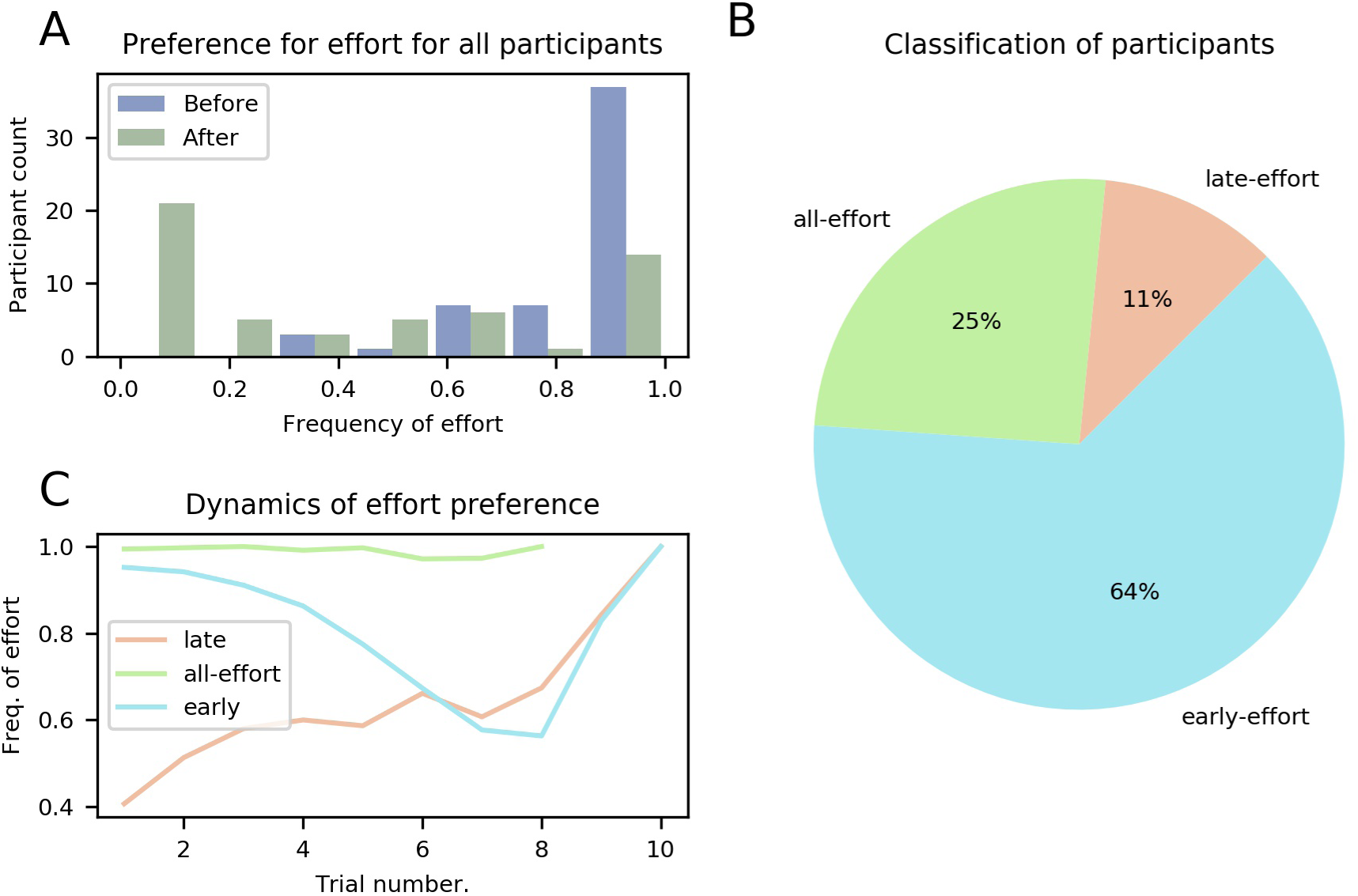
Preference for effort for all participants. **(A)** Histogram of participants’ overall frequency of choosing effort averaged across all trials, separated into before (blue) winning the mini-block and after (green). **(B)** Classification of participants into the three groups all-, early-, and late-effort, see main text. **(C)** Frequency of effort as a function of trial number for the three groups of participants, averaged over participants in each group. Here, only decisions made before the mini-block has been won are included. The different ranges of the lines (e.g. all-effort only reaches trial 8) is due to participants who chose effort more often won the mini-block earlier.

In total, we identified three different groups of participants, differing on when they chose to exert effort (see Figure 2B and Section ‘Dividing participants into groups’ in Methods for more details).

We found that 14 (25%) of all participants continued to choose to do effort even after they had won the mini-block. We refer to these participants as the all-effort group for the rest of this work. In the remaining participants we identified two further distinct categories of behavior when looking at those trials before the mini-block had been won, i.e. trials for which the number of obtained points is smaller than five. The first category comprises six (11%) participants that showed a lower frequency of effortful choices at the beginning of the mini-block, averaged across all mini-blocks, and only later increased their frequency. We refer to these participants as the “late-effort” group. The second category, which included 35 (64%) participants, pertains to participants with the opposite behavior; they started every mini-block with a high frequency of effort and only later in the mini-block, when they had accumulated many points (not necessarily having won the mini-block), started choosing the probabilistic option. We refer to these participants as the “early-effort” group.

We considered that all-effort participants may have misinterpreted the instructions of the task. To discard this possibility, we asked all participants in a post-task questionnaire if they understood that gaining points after filling the bar led to no further reward, to which all participants but one responded that they had understood this; the one participant who responded that she did not understand was part of the all-effort group. Importantly, the task was designed such that all participants could easily win all mini-blocks; we found that across all participants, only four mini-blocks were lost (in all cases by a single point) and no participant lost more than one. We will discuss potential reasons for the choice behavior of the all-effort group in the Discussion.

The model-based analysis results we present in the following sections can account for the all-effort group simply by inferring very low effort-discounting parameters so that the effortful action no longer comes with disutility and thus can be selected freely. However, the choice data of the all-effort group is rather uninformative about the way individuals resolve the dilemma of when to invest effort to reach a goal that is a few trials away, as one might expect given that they always chose to exert effort. Therefore, the all-effort group will be excluded from the following analyses except when explicitly stated.

The dynamics of the frequency with which participants chose the effortful option can be seen in Figure 2C for the three categories of participants (late-, early- and all-effort). For this figure, we averaged, for every trial number, all the choices made by all the participants in each group, using only the trials before the mini-block had been won. It is worth noting that even early-effort participants’ frequency of effort increased in the final two trials. This is because in those mini-blocks when early participants made it to such high trial numbers without having won the mini-block, they urgently needed to accumulate points and thus effort was required to ensure filling the point bar.

### Model-based analysis

In this section, we discuss several hypotheses on how exactly human participants select choices in the sequential task. To do this, we use a series of model-based analyses, using Bayesian model comparison to select the best models.

For all analyses that follow, only trials before the mini-block were used, as only these trials represent goal-seeking behavior.

#### Forward-planning strategies

We first determined which strategy participants used for forward planning, i.e., how they took into consideration all the possible actions that can be taken in the future and their potential outcomes to decide whether they would exert effort or not at any given trial. Effectively, the question we address here is how the discounting models used to describe single-trial behavior are used by participants in tasks that require forward-planning, goal-reaching behavior.

We considered, for each discounting type (effort or probability), two different ways in which participants computed the subjective value of a reward that can only be obtained after several trials. For future efforts, participants may have used either the strategy to apply the effort discount function as many times as necessary to win the mini-block (we call this “stack”), or adding all necessary efforts to win the mini-block and using the discount function on this sum (we call this “add”). For probability, the strategy can be stacking the discount function (“stack”), or calculating the probability of winning by choosing the probabilistic option all remaining trials (“add”). In total, this resulted in four (two variants for effort × two variants for probability; for details see Section ‘Sequential discounting models’). We refer to the model variants as (effort strategy)/(probability strategy), with the four variants being: add/add, add/stack, stack/add, stack/stack. For example, add/stack refers to the strategy where effort is added and probability stacked. To determine which forward-planning strategy was used by participants, we performed formal model comparison between the four forward-planning strategies, following (Stephan et al., 2009).

The results of the model comparison between forward-planning strategies, done by marginalizing over discount functions, can be seen in Figure 3. The posterior distributions over the different variants clearly favor the stack/add variant, with an exceedance probability of ~ 0.99, which means that this forward-planning strategy is orders of magnitude more likely than the others, given the participants’ choices.

**Figure 3.**
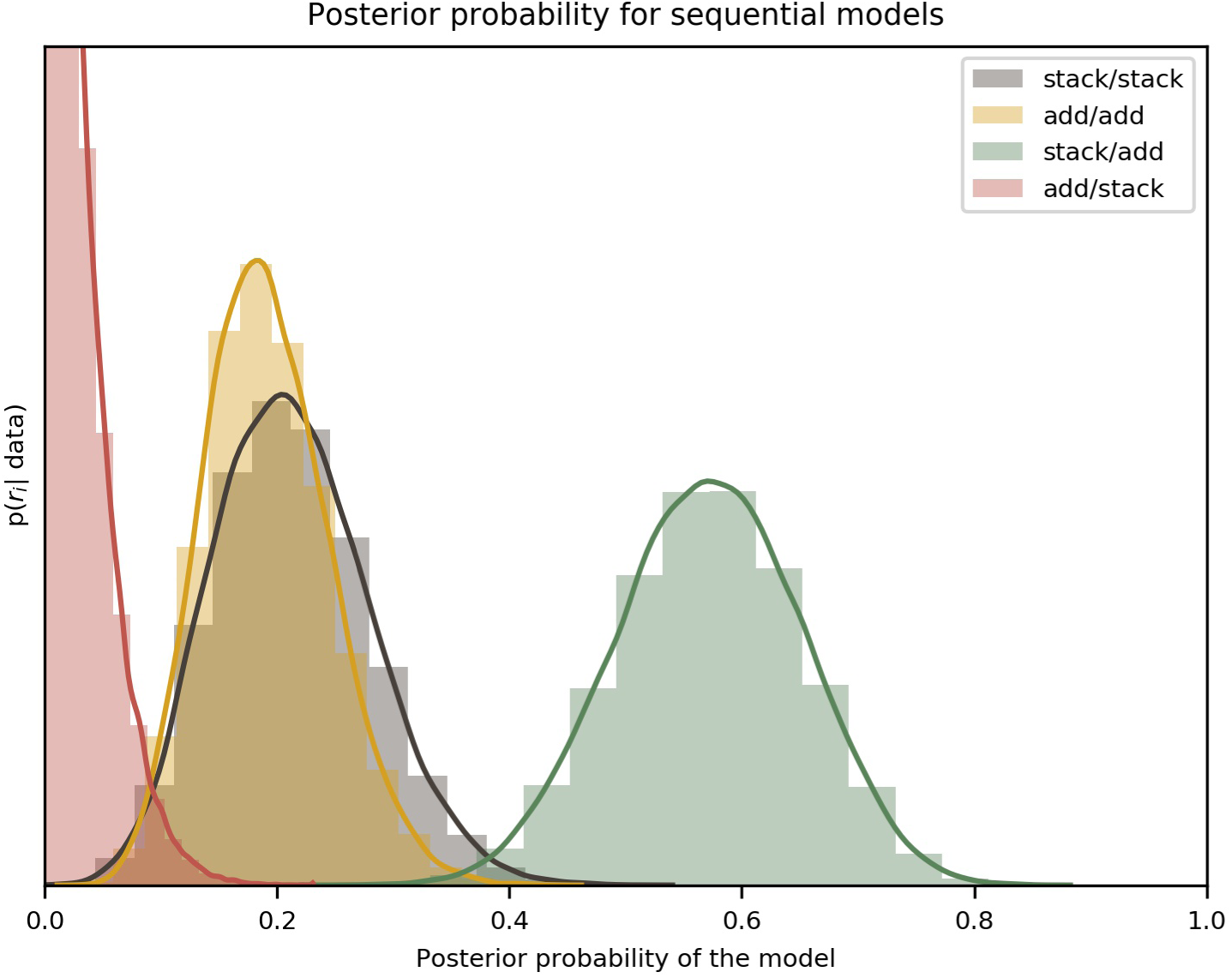
Comparison of the four variants of the sequential model. The label add/stack, for example, refers to effort/probability forward-planning strategy, i.e., effort is added, probability stacked. Each distribution, e.g., green curve and histogram, represents the estimated posterior probability that the model, e.g., stack/add was the best for the data for all participants. The colored lines are an interpolation with a Gaussian kernel. The two effort discount functions (hyperbolic and sigmoid) have been marginalized to compare only the forward-planning components. The y-axis is the probability density of *r*_*i*_ given the data (*p*(*r*_*i*_|data) in Equation 14); the x-axis spans all the possible values of *r*. The peak of the red (add/stack) curve is not shown because the vertical range was cut short for visual clarity.

#### Discount functions

Having selected the forward-planning strategy with the highest posterior probability given the data (i.e. stack/add), we set out to determine which effort discount function (sigmoid or hyperbolic) best fit our participants’ data. To do this, we performed model comparison between the two discount functions. Our results clearly indicate that hyperbolic effort discounting fits the data better than sigmoid discounting, with an exceedance probability ~1.

These analyses were performed with the data of early- and late-effort participants only, excluding the all-effort group. For completeness, we performed the same analysis including all participants and found that the results do not change. This is due to the fact that, for all models, the effort discounting parameter *κ*_*ϵ*_ (from Equation 3) for all-effort participants is estimated to be very low, which causes the model evidence of all models to be the same for that participant. This greatly simplifies model-based data analysis, as it obviates the need for arbitrary exclusion criteria.

#### Modeling effort preferences

Having selected the best-fitting model for the participants’ data (hyperbolic effort discounting, with stack/add forward planning), we show in this section that this model indeed captures participants’ behavior in a measure not directly used for model comparison: the overall frequency of effort for each participant.

To this end, we compared our best-fit models to the experimental data by calculating the overall frequency of effort for each participant across all mini-blocks and doing the same for the models. We performed the analysis only for the early- and late-effort groups. We summarize the results of the comparison in Figure 4A, where we show the observed (experimental) and modeled frequencies of effort for each participant separately. We separated the participants into the late- and early-effort groups; the division is shown as a vertical line, to the left of which are the late-effort and to the right, the early-effort participants.

**Figure 4.**
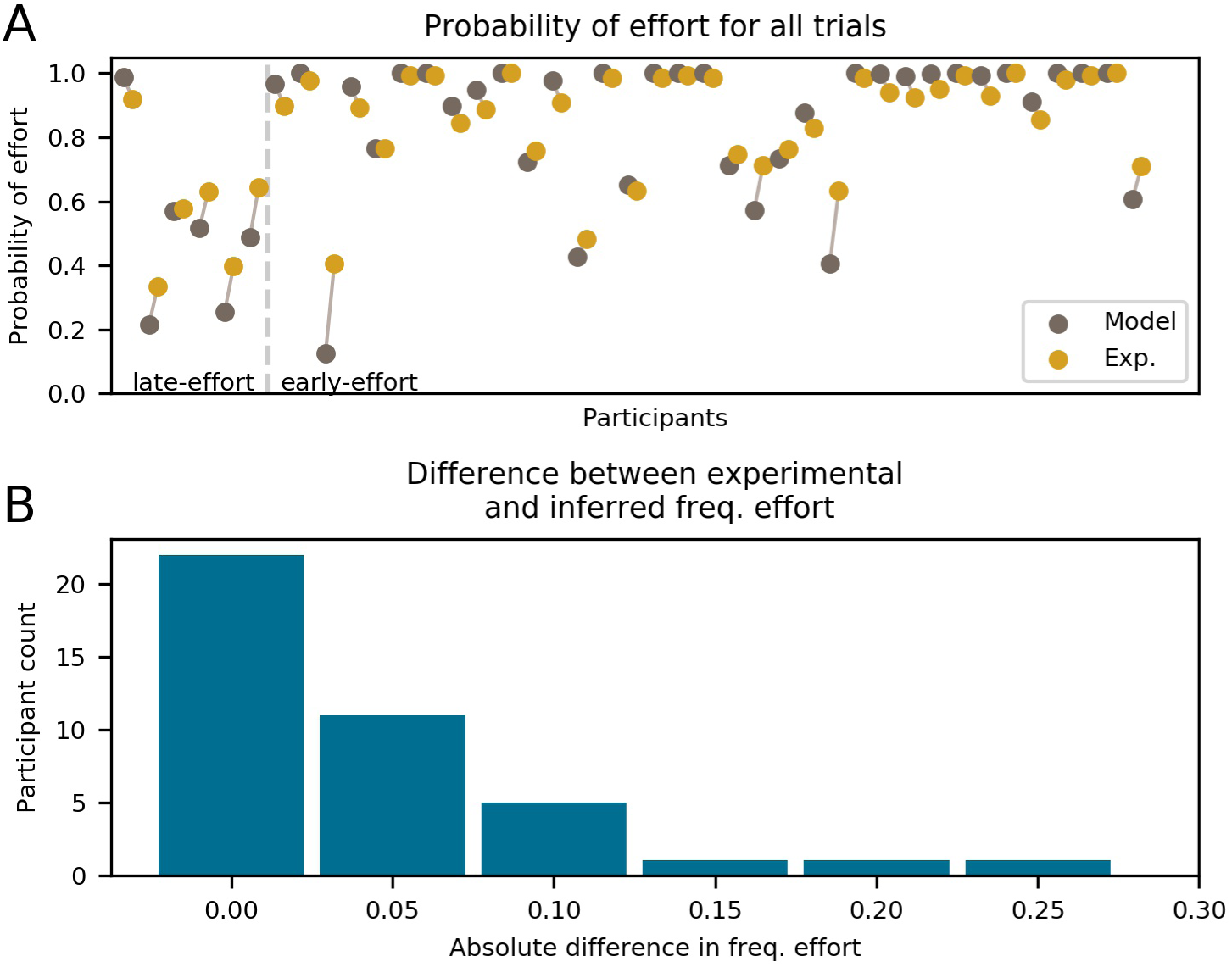
Frequency of effort for each participant (excluding the all-effort group) and the best-fitting models. Only trials before winning the mini-block are included. **(A)** For each participant, two colored dots are shown, which represent the experimental data (green) and the model prediction (brown). Each dot represents the total frequency of effort for the whole experiment. The two dots for each participant are horizontally offset and connected by a line for visual clarity. Participants are divided by the vertical dashed line into late-effort and early-effort. **(B)** Histogram of absolute error between the model and the experimental frequency of error shown in (A).

As can be seen in Figure 4B, the model estimates the probability of choosing effort very well, being within 5% (in frequency of effortful choices) of the experimental data for most participants. Only for three participants we found an error greater than 15%.

It is clear from Figure 4A that the fit is better for higher frequencies of effort. This is due to the fact that the frequency of effort for a participant has lower variability the higher it is, to the point that those participants with an overall frequency of effort (in the early- and late-effort groups) ~ 1 have almost zero variability in their choices.

Note that for the late-effort group in Figure 4A, one participant can be seen with a high frequency of effort. For this participant, effort frequency started very high early in the mini-block and increased as the mini-blocks progressed, meeting our definition of the late-effort group.

#### Effort allocation

In this section, we show that the overall frequency of effort observed in participants can be explained in terms of the discounting parameters fitted from our model. More specifically, we show that participants with a higher frequency of effort are those who discount probability more steeply than effort.

To do this, we selected the best-fitting model, the stack/add, hyperbolic model, and calculated, for each participant, the ratio of the posterior means of the probability discounting parameter *κ*_*p*_ from hyperbolic probability discounting, to *κ*_*ϵ*_ from effort discounting. Figure 5 shows these ratios plotted against the individual overall frequencies of effortful choices. It can be seen that there is a monotonically-increasing relation between the ratio of discount parameters and the overall preference for effort, save for two outliers (one of which has a large absolute difference in Figure 4, belonging to the early-effort group).

**Figure 5.**
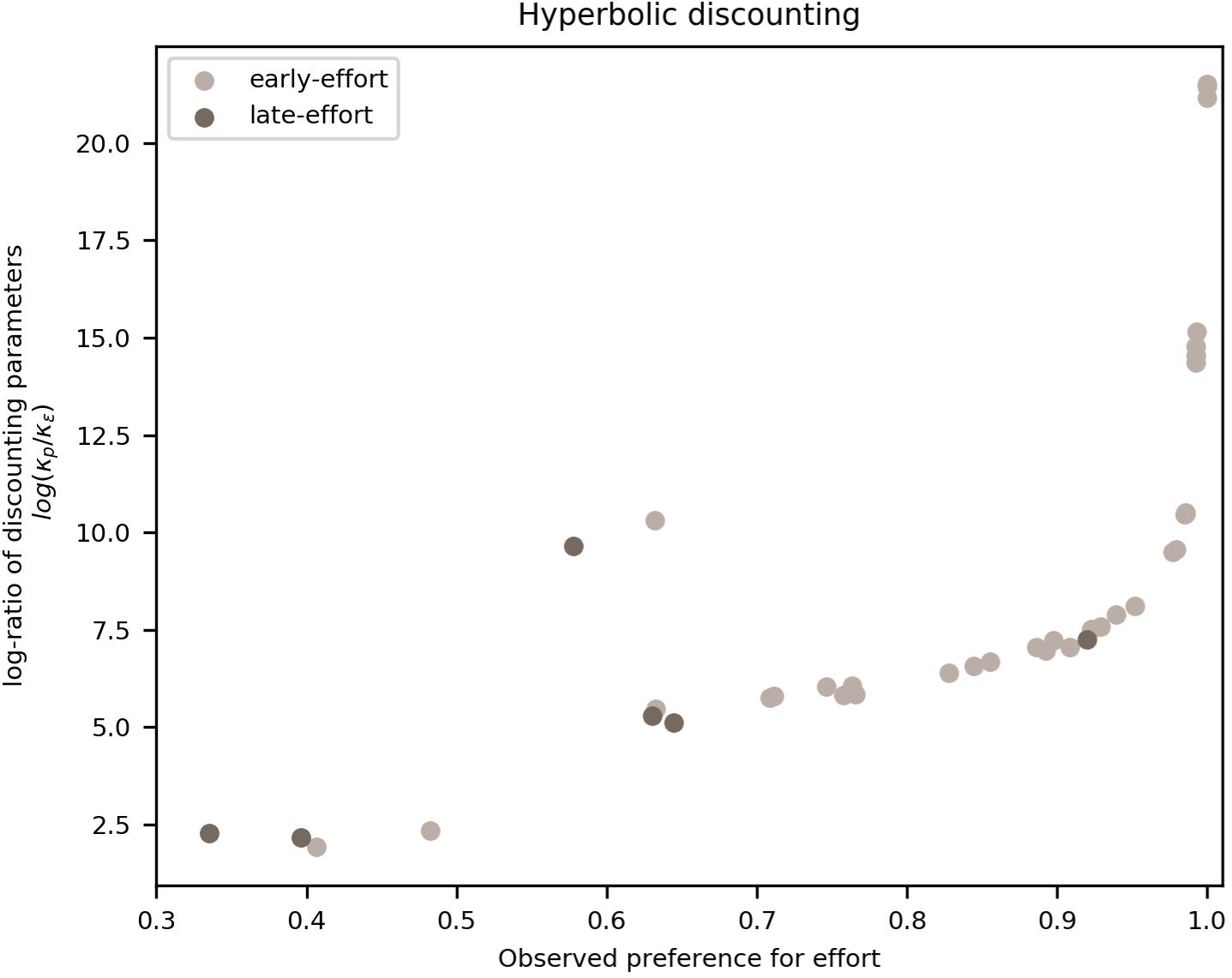
Frequency of effort vs. log-ratio of probability to effort discounting parameters. Each dot represents a participant, divided into late-effort (dark dots) and early-effort (light dots). We plot the frequency with which a participant chose the effortful action (until reaching the goal of a mini-block) on the x-axis and the log-ratio of the parameters for probability to effort discounting, i.e. *κ*_*p*_ to *κ*_*ϵ*_, on the y-axis (log-scale for clarity).

This monotonically-increasing relation can be interpreted in terms of the comparison between the two options in the task: a participant with a high ratio discounts probability more steeply than effort, which translates into a lower valuation of any probabilistic offer, compared to an effortful one. At values of the frequency of effort ~1, the log-ratio increases rapidly (faster than exponentially) due to the nature of the model, as the probability of effort grows more slowly than exponentially as *κ*_*ϵ*_ decreases linearly.

## Discussion

We designed a sequential decision-making task in which participants could choose, in each trial, to exert mental effort in order to improve their chances of obtaining reward at the end of a mini-block (i.e., sequence) of ten trials. In this task, participants had the option to exert effort immediately to ensure future reward or choose a probabilistic option and wait until a later trial to re-evaluate if effort needed to be exerted. With this task, we aimed at determining when participants choose to exert effort and which forward-planning strategy they employ to make such a decision. To this end, we proposed a forward-planning model for goal-directed, sequential decision-making behavior that incorporates different strategies for the consideration of future exertion of effort.

Our results show inter-participant variation in when they chose to exert effort, with most participants choosing to start a mini-block with effort and only later chose to not exert effort. Additionally, the results of our model comparison between four different forward-planning strategies show that most participants considered future efforts by stacking the effort discount function, i.e., by applying the function as many times as they planned to exert effort in future trials. For probability discounting, we found that the best-fitting model calculates the overall probability of reaching the goal (winning a mini-block) when always choosing the probabilistic option. We also found that hyperbolic effort discounting fits the data of our experiment better than sigmoid effort discounting. Finally, we showed that the overall frequency of effort for a participant can be explained by the ratio of the inferred probability discounting to the effort discounting parameters.

### Preference for effort

We found that most participants had a strong preference for effort. A quarter of participants (the all-effort group) went as far as choosing to exert effort even when it brought no extra monetary reward. In particular, participants in the all-effort group did not seem to be following the instructions of the task. A similar phenomenon, i.e., continuing to exert effort when it no longer is necessary, has been observed in physical effort experiments (Bouc et al., 2016; Schmidt et al., 2008).

There may be two possible reasons for this phenomenon: First, the level of cognitive effort in our number-sorting task was probably not high enough to trigger a cost/benefit analysis in participants in the all-effort group. In our task, the effortful option came implicitly tied to an increase in the probability of earning monetary reward, which added to the overall benefit of exerting some effort. Moreover, other reasons may be that for some participants, the number-sorting task was interesting on its own (Inzlicht, Shenhav, & Olivola, 2018), participants did not want to wait for the next trial while doing nothing, and wanted to make sure they did not lose practice, all of which were reported by our participants in a post-task questionnaire. A related possibility was suggested by Pessiglione et al. (2018), namely that participants might want to “make an impression on the experimenter” by always choosing to exert effort.

Second, we speculate that highly motivated individuals might “flatten” their effort discount curves (e.g., by making *κ*_*ϵ*_ smaller) to more easily attain highly-valued rewards in a scenrario like a psychological experiment, which they might misunderstand as a competetive scenario. This context-dependent meta-control could come in the form of a meta-parameter that controls the size of *κ*_*ϵ*_ depending on higher-level goals (e.g. “how much do I want to win each mini-block?”). As volunteer participants can be assumed to be highly motivated, especially when monetary reward is contingent on performance (Hertwig & Ortmann, 2001), this would mean that their effort discounting parameters are lower, causing the observed high frequency of effort.

Testing these two possible explanations could prove fruitful in future research. Testing the low-effort level possibility would require a task that parametrically varies the effort level to establish higher levels of cognitive effort, as is done typically with physical effort (Prévost et al., 2010). Based on these variations, the proposed model-based approach can be used to infer meta-control by establishing differences in individual effort and probability discounting parameters between different levels of effort requirements.

### Action sequences

As part of the present model’s definition, we limited the action sequences considered by the model to the all-effort (*π*_*ϵ*_) and the all-probability (*π*_*p*_) action sequences (see Section ‘Sequential discounting models’). Here, we discuss the reasoning behind this choice and its ramifications.

We posit that as a means to prune the decision tree, participants developed a strategy in which they evaluate the current state of the task and determine it to be “good” or “bad”, which in turn allowed them to simplify the decision tree to the two action sequences *π*_*ϵ*_ and *π*_*p*_. A good state is one in which the participant is close to winning. A bad state is one in which losing seems likely. A good state is then one in which the participant can afford to choose the probabilistic option without it becoming too likely to lose the mini-block, while a bad one is one in which effort needs to be exerted to continue to have a chance at winning. It depends on the participant where exactly this change from good to bad state lies.

In a bad state, effort is, by definition of the bad state, necessary not only in the current trial, but also for all the remaining ones, as otherwise the probabilistic option would still be viable and the state would be good. Therefore considering a mixed action sequence (i.e. one in which both effort and probability can be planned for future trials) is unnecessary in bad states.

In contrast, in a good state, the probabilistic option is still viable. This definition does not preclude future necessity of effort, as things could go wrong and all probabilistic options be lost, which eventually would lead to a bad state. However, as states are evaluated at every trial during the experiment, it is unnecessary to consider this possibility when evaluating the action sequences during a good state; instead, the participant can simply wait until the state has actually become bad in the future and then switch to the all-effort strategy. This implies that good states only require the evaluation of *π*_*p*_.

How is this state evaluation carried out? Since the only viable option in a good state is *π*_*p*_ and the only viable option in a bad state is *π*_*ϵ*_, one can turn this around and define a good state as one in which *z*(*π*_*p*_) > *z*(*π*_*ϵ*_), where *z*(·) is the valuation function (Equation 7), and a bad state as one in which the opposite is true. Therefore, the decision-making agent can decide between effort and probability by comparing the valuations of *π*_*p*_ and *π*_*ϵ*_, as done in the proposed model. This evaluation could be affected by the meta-control we discussed in Section ‘Preference for effort’; for example, a highly-motivated individual would classify states as “bad” more often than one with low motivation. Whether motivation and, more generally, meta-control could change which action sequences are evaluated at all should be the target of future research.

This evaluation could be affected by the meta-control we discussed in Section ‘Preference for effort’; for example, a highly-motivated individual would classify states as “bad” more often than one with low motivation. Whether motivation and, more generally, meta-control could change which action sequences are evaluated at all should be the target of future research.

### Effort and goal reaching

It has been suggested that individuals generally tend to avoid cognitive effort (Kool et al., 2010; Westbrook et al., 2013). However, in the tasks used in the experiments by Kool et al. (2010) and Westbrook et al. (2013), there was no set goal that could be reached more readily via the exertion of cognitive effort. In the study by Kool et al. (2010), participants could not earn additional money if they chose the more effortful task more often. In the experiments in (Westbrook et al., 2013), the association between the actual investment of effort in an increasingly difficult *n*-back task, the choice behavior in the titration procedure used to determine the subjective value of redoing the different *n*-back levels, and the actual payment based on four randomly selected choices in the titration procedure may simply have been too unconstrained. In the present task, it was clear in every trial and mini-block that choosing the effortful option would be beneficial for obtaining the reward.

In the present experiment, the frequency of choices of the effortful option was lower after a mini-block had been won (see Figure 2A). The early-effort group tried to reduce effort during the course of a mini-block and increased it only towards the end of the mini-block if they had not already won that block (see Figure 2B). Likewise, the late-effort group only gradually increased their effort, especially in later trials of a mini-block when facing to lose the mini-block. Thus, our results suggest that individuals may tend to avoid cognitive effort unless its exertion is necessary to reach their goals.

This caveat to the assumption of a general tendency of individuals to avoid the exertion of cognitive effort is also backed by the observation that stable individual differences in personality traits related to the tendency to willingly exert cognitive effort have been found to be associated with effort discounting: Kool and Botvinick (2013) found that individuals with higher scores in Self-Control showed less avoidance of cognitive demand, and Westbrook et al. (2013) observed that participants with higher scores in Need for Cognition showed less effort discounting. While Self-Control is characterized by the investment of mental effort to control one’s impulses that interfere with long-term goals (Tangney, Baumeister, & Boone, 2004), Need for Cognition refers to the tendency to engage in and enjoy effortful mental activities (Cacioppo, Petty, Feinstein, & Jarvis, 1996), which can be summarized as cognitive motivation. It remains to be determined whether our participants’ habitual cognitive motivation may have played a modulatory role in their decisions to choose the effortful condition more frequently because of their intrinsic motivation to invest cognitive effort. Taken together, our results partly corroborate the seminal findings by Kool and Botvinick (2013) and Westbrook et al. (2013) in pointing to individual differences in the willingness to invest cognitive effort and extend them by showing that the assumption of a general tendency for the avoidance of the exertion of cognitive effort only holds if there is no goal to be attained that can be achieved more readily by the exertion of effort.

Our computational modelling approach shares similarities with three other areas of research. First, to model goal reaching in sequential decision making tasks with probabilistic outcomes, some method of forward planning based on computing and evaluating the consequences of action sequences is required, e.g. using the active inference framework (Cuevas Rivera, Ott, Markovic, Strobel, & Kiebel, 2018; FitzGerald, Schwartenbeck, Moutoussis, Dolan, & Friston, 2015; Schwartenbeck, FitzGerald, Mathys, Dolan, & Friston, 2015) or – as in the present study – using backward induction based on maximization of expected value (e.g., Kolling, Wittmann, & Rushworth, 2014; Korn & Bach, 2018). We add to this modelling work by showing how one can compute the apparent cost of exerting mental effort to reach a goal after several trials.

Second, modelling the dilemma between the apparent costs of exerting effort and reaching a goal with high probability is a long-standing research question at the interface between psychology and economics. For example, Steel and König (2006) describe a computational framework that aims at qualitatively modelling, as a function of time towards a deadline, how people compute preferences of options. Their model of the procrastination phenomenon, based on a preference reversal close to the goal, is similar to the proposed mechanism how the late-effort group increases their effort frequency close to the goal. The main difference of the here-proposed approach and the approach taken by Steel and König (2006) is that we use probabilistic inference and model comparison to fit several models to participant data and select the best among alternative models.

Third, it is still an open question whether individuals with a lower tolerance for risk should be more willing to exert effort to increase their chances of winning, e.g., (Briys & Schlesinger, 1990; Jullien, Salanié, & Salanié, 1999). However, while these studies presented a mathematical framework to compare effort and probability discounting, they did not allow for the evolution of the state of the task, making it more akin to single-trial discounting paradigms and models.

The model we present is highly generalizable by modifying the underlying transition structure. This makes the model applicable to any sequential decision-making task which involves the exertion of effort, as well as considerations of probability and delays. This can enable researchers to characterize, using the model parameters, forward-planning, goal-reaching behavior in many tasks, for example in foraging tasks, where plans for future patch switches can incorporate the associated costs in effort (Kolling, Behrens, Mars, & Rushworth, 2012; Shenhav, Straccia, Cohen, & Botvinick, 2014, e.g.). Moreover, more complex effort/reward structures can be modeled, such as sub-goals necessary for the main goal, or states that open/close doors to reward options.

In conclusion, we have presented a novel combination of a sequential decision making task and a computational model based on discounting effects to describe how participants plan forward to exert effort to reach a goal. We believe that this computational-experimental approach will be highly useful for future studies in the analysis of how participants meta-control the cost/benefit ratio during goal reaching.

